# Livestock as vectors of organic matter and nutrient loading in aquatic ecosystems in African savannas

**DOI:** 10.1101/2021.07.13.452213

**Authors:** Jacob O. Iteba, Thomas Hein, Gabriel A. Singer, Frank O. Masese

## Abstract

Populations of large wildlife have declined in many landscapes around the world, and have been replaced or displaced by livestock. The consequences of these changes on the transfer of organic matter (OM) and nutrients from terrestrial to aquatic ecosystems are not well understood. We used behavioural data, excretion and egestion rates and C: N: P stoichiometry of dung and urine of zebu cattle, to develop a metabolism-based estimate of loading rates of OM (dung), C, N and P into the Mara River, Kenya. We also directly measured the deposition of OM and urine by cattle into the river during watering. Per head, zebu cattle excrete and/or egest 25.6 g dry matter (DM, 99.6 g wet mass; metabolism) - 27.7 g DM (direct input) of OM, 16.0-21.8 g C, 5.9-9.6 g N, and 0.3-0.5 g P per day into the river. To replace loading rates OM of an individual hippopotamus by cattle, around 100 individuals will be needed, but much less for different elements. In parts of the investigated sub-catchments loading rates by cattle were equivalent to or higher than that of the hippopotamus. The patterns of increased suspended materials and nutrients as a result of livestock activity fit into historical findings on nutrients concentrations, dissolved organic carbon and other variables in agricultural and livestock areas in the Mara River basin. Changing these patterns of OM and nutrients transport and cycling are having significant effects on the structure and functioning of both terrestrial and aquatic ecosystems.

## Introduction

Large animals strengthen the linkage between ecosystems by facilitating the movement of organic matter and inorganic nutrients, often against naturally-established boundaries (1, 2). For instance, when animals spend time in a recipient ecosystem after feeding elsewhere, they directly contribute carbon and nutrients to that ecosystem through excretion and egestion (3–5). Similarly, the death of animals can represent a material flux between ecosystems (6, 7). One of the greatest examples of subsidy transfer by mammals through carcasses includes “whale falls” when dead whales sink to the seafloor resulting in an enormous loading of pulsed organic matter and nutrients (8). Another example is the nearly annual mass drowning of wildebeest (*Connochaetes taurinus*) in the Mara River, East Africa during the Serengeti-Maasai Mara migrations (7).

For landscapes hosting huge populations of large mammalian herbivores (LMH), transfer of organic matter and nutrients from terrestrial to aquatic ecosystems has been a subject of great research interest (9–11). These inputs are often judged as negative for water quality, biodiversity and ecosystem functioning. For instance, increased input of cattle dung into streams and rivers can cause microbial contamination and eutrophication (12–14). Livestock activity can also mobilize sediments which, in addition to the fine particulates in excreta, can increase turbidity in the aquatic ecosystems, which may reduce light penetration and limit primary production. Similar to livestock, increased turbidity in rivers has also been linked with the presence of hippos, which also have high levels of loading of organic matter and nutrients (4, 15), which have been linked with poor water quality, hypoxia, loss of fish and invertebrate diversity, and altered ecosystem functioning (12, 15, 16). However, terrigenous materials by native LMH are vital subsidies driving the natural structure and function of riverine ecosystems draining savanna grasslands and grazing areas (7, 17, 18). Consequentially, declining populations of wild LMH in many regions around the world and their replacement by livestock (19–21) raises questions on the ecological consequences of such a replacement on the structure and functioning of aquatic ecosystems (22).

Similar to wild LMH such as hippopotamus, cattle are mobile consumers capable of moving resources from savanna grasslands to aquatic systems (4). In addition to direct input by defecation and urination during watering or crossing (23), attached faeces washes from cattle feet and disturbance of sediment re-suspends material into the water column (24, 25). Further, livestock can facilitate subsidy transfer by the promotion of soil and riverbank erosion (11). In how far livestock can replace wildlife as a vector of terrestrial subsidies depends on the similarity of the subsidy in terms of quantity, quality and timing and duration (5, 18, 22). These are influenced by several species-specific factors, including body size, population size and behaviour linked to water (i.e., ontogenetic habitat switch, migration, feeding) (5, 10). Water-dependent grazers that are obligate drinkers have the potential to transfer more subsidies than water-independent browsers that visit watering points only occasionally (26). For livestock, management decisions determine the timing and duration of interactions with aquatic environments. For instance, paddocking or fencing and herding restrict access to watering points and, hence, the possibility of egestion or excretion in aquatic ecosystems (23, 27). In contrast, unrestricted livestock access to watercourses creates footpaths where nutrients and organic matter are connected to waterbodies through hydrologic vectors (28, 29).

Several studies have quantified inputs of organic matter (dung) and nutrients by either wild LMH or livestock to disparate aquatic ecosystems (4, 9, 10, 12). For African savannas, available data for some wild LMH (4, 10, 30) contrasts the lack of comparative data for livestock, even though livestock graze side by side with or have completely replaced wildlife (20, 31–33). Whether livestock can quantitatively and qualitatively replace wild LMH as vectors of terrestrial subsidies to aquatic ecosystems is unknown (22). Thus, data-driven models on nutrient balances in both grazing and farming systems (34) are required to understand the implications of growing livestock populations on water quality and ecosystem structure and functioning of streams and rivers.

Here, we quantified loading rates of organic matter and nutrients by cattle into an African savanna river, that supports large populations of both livestock and wild LMH (20, 33, 35). The objectives were to 1) quantify livestock-mediated subsidies by assessing behaviour in concert with excretion and egestion across sites with varying densities of cattle, 2) compare these data with previously reported inputs by hippos (4), and 3) determine the influence of livestock access (watering points) on water quality.

## Methods

The research permit for conducting this study was granted by the National Council For Science, Technology & Innovation, Kenya. The methods for calculating loading rates of organic matter (OM) and nutrients by livestock and hippos have been borrowed from (4) (2015) and (22) (2020). However, the (22) (2020) paper only has estimates for organic matter (dung) for livestock, and here we present data on OM, carbon (C), nitrogen (N) and phosphorus (P) for both urine and dung. This study also extrapolates the loading estimates for river-reaches to the catchment scale.

### Study area

This study was conducted in the Mara River (MR) basin, Kenya/Tanzania. The Mara River has its source in the Mau Escarpment in Kenya and drains into Lake Victoria in Tanzania. As the only perennial river, the Mara River is very important for watering wildlife migrating between the Serengeti National Park (SNP) in Tanzania and the Maasai Mara National Reserve (MMNR) in Kenya during the dry season (36). Extensive grasslands in the pastoral areas adjacent to the river also provide dispersal ranges for resident wildlife (20, 33, 37).

In the recent years, the declining wildlife numbers in the SNP-MMNR ecosystem have been linked to the intensification of land use, expansion of agriculture, sedentarization of once pastoral communities and diversification of livelihoods (20, 38, 39). The decline in wildlife numbers is paralleled by growing populations of livestock intruding into protected areas (33, 40). The biomass of livestock as a per cent of total livestock and wildlife biomass recorded within the MMNR boundaries has increased from an average of 2% in the 1970s to 23% in the 2000s; over the last decade, livestock biomass has become more than 8 times greater than that of any resident wildlife species (20).

In the Middle Mara and Talek regions, livestock numbers are significantly higher than the rest of the MR basin (38). These regions are also home to Maasai pastoralists who graze over 200,000 cattle and higher numbers of sheep and goats in communal lands adjoining the MMNR and utilize streams and rivers as watering points and crossings (38). In the communal conservancies outside the MMNR, people graze their livestock in a manner that allows livestock to co-exist with wildlife (20, 33, 35). This results in a spatial pattern with hippopotamus inside the MMNR, mixed hippo and livestock (cattle, goats and sheep) present in areas adjoining the MMNR and only livestock grazing areas further away from the MMNR and conservancies. This spatial distribution reflects the ongoing replacement of native wildlife with essentially exotic livestock.

### Study design

The MR basin was divided into 5 regions defined by elevation, catchment land use and livestock densities; Nyangores, Amala, Middle Mara and Talek River and MMNR (Figure 1). Sites were selected at livestock watering points in each of the five regions for livestock (cattle, goats and sheep) census, observation of behaviour and periodicity of interactions with streams and rivers during the dry season in February-March 2017. Because of logistical constraints only the Talek Region sites were monitored and sampled for reach-scale effects of livestock access on water quality and nutrient concentrations during the dry and wet season in November-December 2017. Figure S1 (supplementary information) provides context for the discharge of the major rivers during the time of sampling. Sites in the Nyangores and Amala regions were located in areas with low to medium densities of livestock (<50 individuals per km^2^) (41) as most of the inhabitants in these regions are also involved in smallholder mixed agriculture (livestock rearing, cash and subsistence crops such as tea, maize and potatoes). Sites in the Middle Mara and Talek regions had higher livestock densities (average of >100 individuals per km^2^) (41) with over 250,000 cattle present all year round (20, 33, 38, 42). The Middle Mara, Talek and MMNR regions also host over 4000 hippopotami (43). In total 66 sites were selected for the study: 21 sites in the Nyangores, 17 sites in the Amala, 7 sites in the Middle Mara, 16 sites in the Talek and 5 sites in the MMNR.

**Figure 1:**
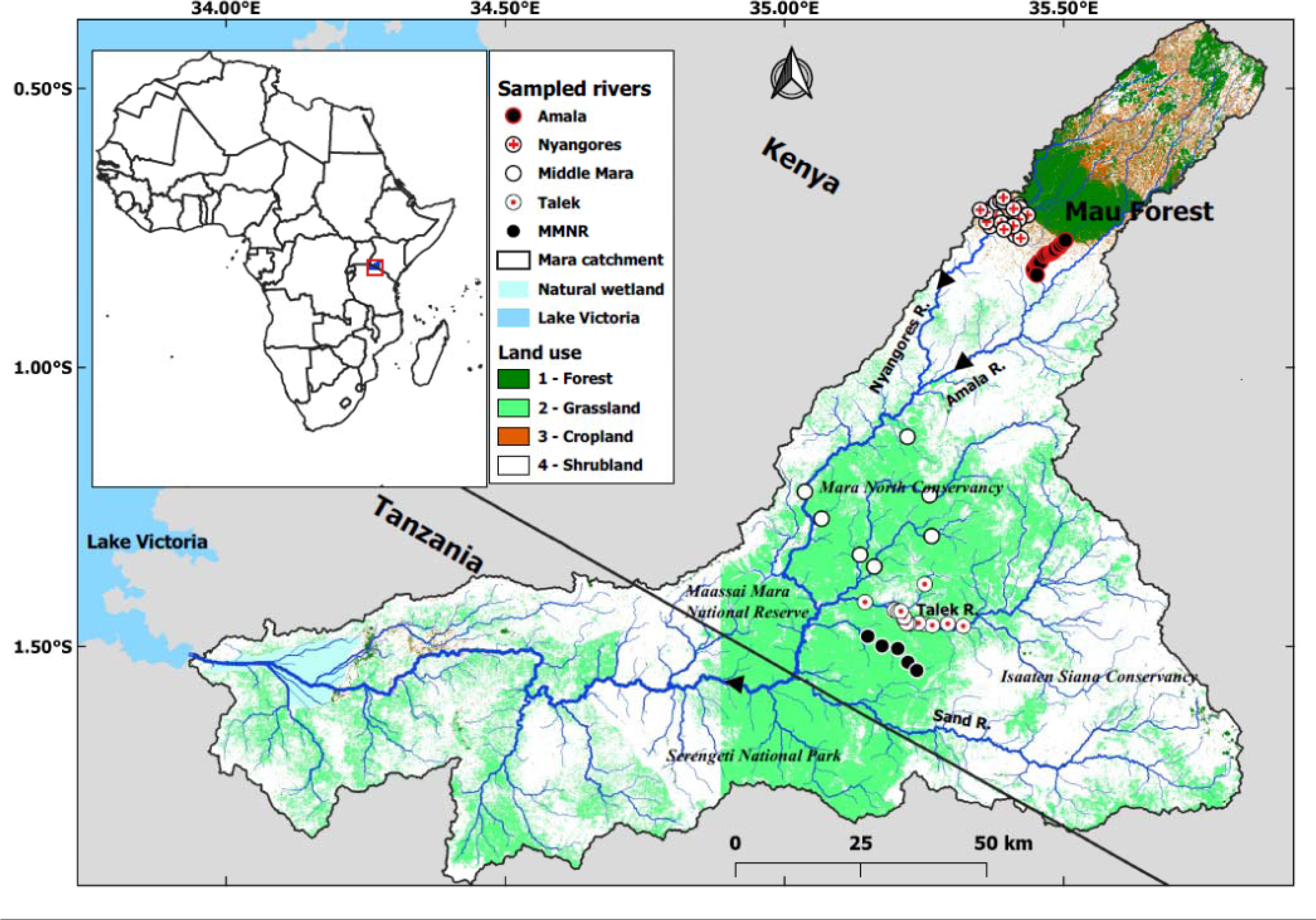
Map of the study area showing the location of the livestock study sites in the four regions in the Mara River Basin, Kenya. The MMNR sites are within the Maasai Mara National Reserve.

### Sampling methods for water quality

Water samples were collected immediately upstream and downstream of livestock watering points in the Talek region during the dry and wet seasons. A portable meter (556 MPS, Yellow Springs Instruments, Ohio, USA) was used for measuring temperature, dissolved oxygen concentration, electrical conductivity and pH *in situ*. Known volumes of river water were directly filtered (GF/F) into acid-washed HDPE bottles for analysis of nutrients and dissolved organic carbon (DOC) concentrations. DOC samples were acidified to pH <2 before further preservation. Replicate filters were used for the measurement of water column chlorophyll-a, total suspended solids (TSS) and particulate organic matter (POM). Sediment samples were collected using corers (diameter 10 cm) and placed in aluminium envelopes for analysis of organic matter and nutrients. For benthic chlorophyll-a analysis, a known area of the stone substrate was scraped off and the slurry was then filtered through GF/F filters. All water and sediment samples were kept at 4 °C during transport to the laboratory where they were either analyzed immediately or frozen until analysis. All chlorophyll-a samples were wrapped in aluminium foil, transported using a cooler box with ice, and stored frozen in the laboratory pending analysis. For *in situ* measured variables and nutrients, sampling was done thrice a day (morning, noon and evening) to capture diel variation in numbers of cattle. The mean differences in physico-chemical variable and nutrients between upstream and downstream reaches of watering points and between morning (no livestock), noon (increased livestock numbers) and evening (reduced livestock numbers) were used to assess the effects of cattle for selected watering points.

### Livestock behaviour and direct loading estimation at watering points

At the observation sites in the Nyangores, Amala and Talek regions, we assessed numbers and behaviour of livestock during the day from 9:00 am to 18:00 h on a random day in the dry and wet seasons. We recorded the number of livestock visiting a watering point, the number defecating and/or urinating in or near the river (not all cattle visiting a watering point or site do so), and the time spent in or near the river. Often enumerators would stand at a safe distance 10-20 m away from the stream on a raised ground to see all the livestock in the water. In addition to recording livestock behaviour using a questionnaire, photos and short videos were used to analyse livestock behaviour and activity for later verification (Figure 2). Because of large numbers of livestock visiting a watering point in the Talek region, observations were done by two people per site. Despite the large numbers of livestock, watering was often done in shifts as not all cattle could drink water at the same time. Moreover, individual herders arrived at watering points at different times during the day, and this gave enumerators ample time to count and monitor instream livestock activity. Because of the low number of cattle visiting watering points in the Nyangores and Amala regions, one person was able to count and monitor livestock bahaviour and activity at the watering points.

**Figure 2.**
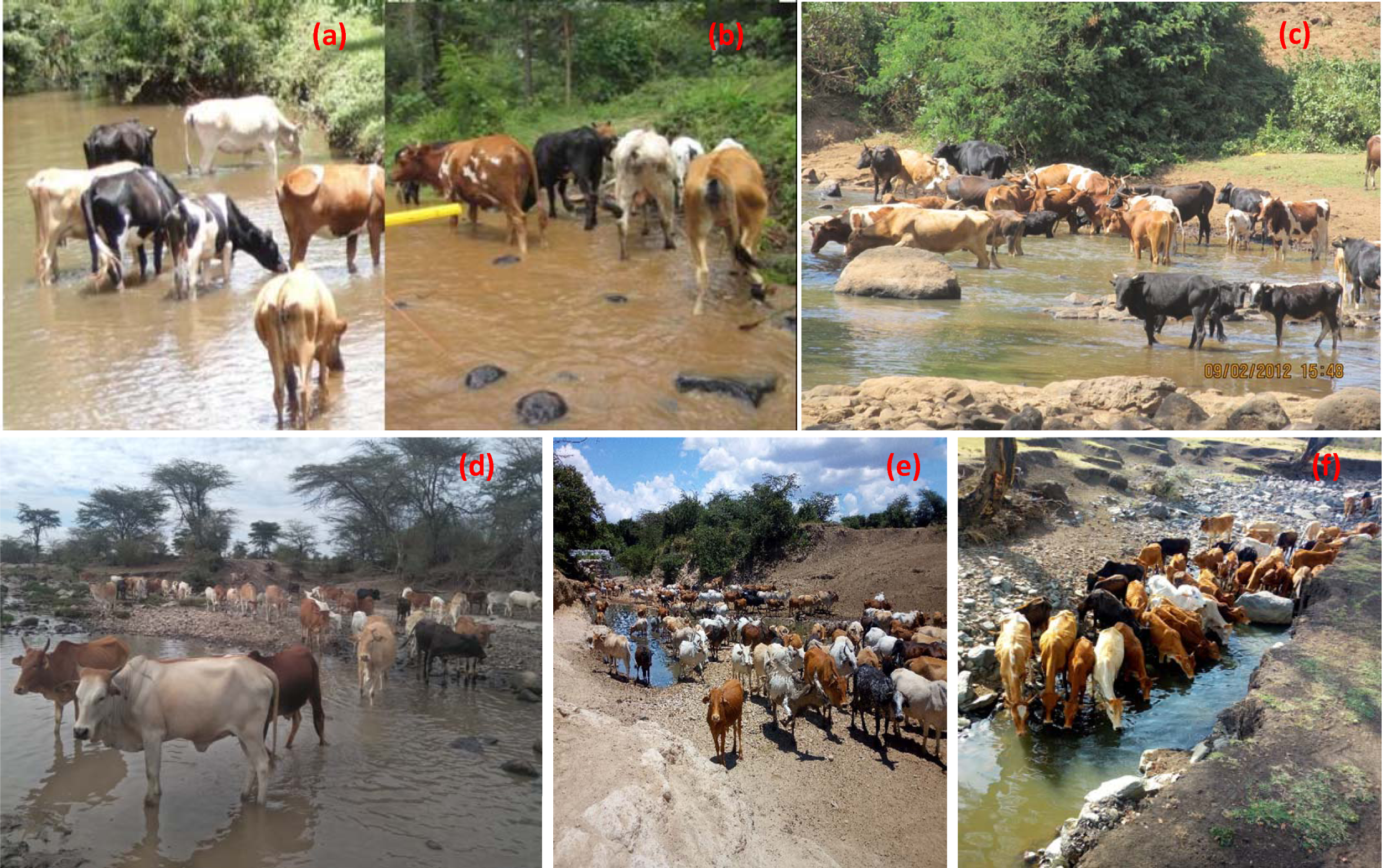
Livestock watering points during monitoring of behaviour in the upper Mara River Basin (a, b and c) and lower basin in the Talek Region (d, e and f) during the wet (a, b and c) and dry (c, e and f) seasons.

At each site, fresh cattle dung from individual on-shore defecation of both adult and sub-adult cattle were weighed per defecation event. Subsamples of dung were collected for wet-dry weight conversion and analysis of C:N:P stoichiometry. Because of logistical constraints, it was not possible to directly measure the volume of urine produced, but we collected urine samples for measurement of C:N:P stoichiometry. Urine was collected from livestock early in the morning in bomas before going out for grazing. The collection was done manually by holding a container against trickling urine from individual cattle. The average urine volume per urination event was estimated as 0.66 L (see below). The following equation determined the amount of nutrients (C, N and P) and organic matter input per cattle per day:

Mass of excretion/egestion_C,N,P_ = Weight of dung/urine Х Content of dung/urine_C,N,P_

The average per capita deposition of faeces and urine directly into or near the river were computed by multiplying average faeces weight or urine volume with the proportion of actually defecating or urinating individuals. In this study we noted that cattle visit the river at least once per day, and for loading estimates we only used single visits per cattle head per day. This decision is based on our livestock movement and herding behavioural data. For instance, in the upper Nyangores and Amala region livestock rearing is done in paddocks, and farmers lead their cattle to watering points once a day, usually around noon to early evening, and return them to the paddocks until the following day. Similar behaviour was noted in the lower Mara River basin (Talek and MMNR) where herders mostly drove their livestock to watering points around mid-day hours and returned them back to grazing grounds far from watering points.

### Indirect loading estimation based on a metabolism model

In addition to direct loading measurements, we developed a simple metabolic model to estimate cattle loading rates of organic matter (dung) and nutrients (C, N and P) from dung and urine deposited by cattle into the Mara River (see Supplementary Information 1), and compared results with existing estimates of loading rates for hippos in the river (4). We estimated cattle loading rates of OM, C, N and P as a fraction of daily dry matter intake (DMI), the proportion of dung (organic matter, OM) egested or excreted, the volume of urine produced and time spent in the river, and we multiplied the per-cattle loading rate by the cattle population to get the total loading rates for all cattle. We used the average stoichiometry of cattle faeces and urine for each region to determine the loading rates of C, N and P from egestion and excretion. We then compared the loading of cattle and hippopotamus dung in areas of the Mara River where their distributions overlap.

### Upscaling loading to region-wide subsidy fluxes and comparison with wild LMH

Livestock census data were obtained from the National and County Ministries of Agriculture and literature to determine densities resident in each of the regions studied. Assuming that all cattle visit the river only once per day, we then estimated total loading rates in the five regions by multiplying the per-capita loading rate with cattle population and compared these with the hippopotamus population in the three regions where their distribution overlap (Middle Mara River, Talek River and MMNR). Loading estimates for dung and urine were translated and integrated to total subsidy fluxes of C, N and P using their respective stoichiometries (see below). Data on livestock were obtained from the Ministry of Agriculture and Kenya National Bureau of Statistics reports (44–47). Livestock and wildlife (hippos) data were also obtained from unpublished and published survey reports (20, 33, 38, 42, 43, 48).

### Laboratory analyses

Water samples: Dissolved nutrient fractions including total dissolved nitrogen (TDN), soluble reactive phosphorus (SRP), nitrates (NO_3_^-^), and ammonium (NH_4_ ^+^) were analysed from filtered water samples, while unfiltered water was used for total phosphorus (TP) and total nitrogen (TN) analysis [24]. TP, TN and SRP were determined using standard colourimetric methods. NO_3_ ^-^was analysed using the salicylate method and NH_4_^+^ was analysed using the reaction between sodium salicylate and hypochlorite solutions. Dissolved organic carbon (DOC) and total dissolved nitrogen (TDN) concentrations were determined using a Shimadzu TOC-V-CPN. TSS and POM (as ash-free dry mass after combustion at 450°C for 4 hours) were determined gravimetrically (49). Chlorophyll-a was extracted from the GF/F filters using 90% acetone solution and assessed spectrophotometrically at the University of Eldoret (49).

#### Nutrients in sediments

TN and TP were determined colourimetrically after acid digestion of oven-dried samples. Colourimetric procedures were applied for the analysis of NO_3_^−^ and NH_4_ ^+^ from wet sediments after extraction using 0.5M K_2_SO_4_. Inorganic phosphorous concentration was determined using extraction after Olsen with 0.5 M sodium bicarbonate at pH 8.5 (50).

#### Stoichiometry of dung and urine

Dung samples were analysed for C, N and P content. For C and N, dried (60⁰C for 48h) samples were grounded, weighed and loaded into tin cups, and analysed on an elemental analyser (Hekatech-Elemental analyser, Thermo Finnigan). For P, samples were weighed, ashed in a muffle furnace at 550 °C and analyzed following the persulfate digestion method (51). Because of logistical constraints, samples were collected from the Talek region only for analysis of C, N and P in the urine. The urine samples were analysed for total dissolved nitrogen (TDN), total dissolved phosphorus (TDP) and dissolved organic carbon (DOC).

### Data analysis

Non-parametric, rank-based H-tests (Kruskal-Wallis ANOVAs) was used to test for differences in stream size (width, depth and discharge) at the various watering points (sampling sites) in the Amala, Nyangores and Talek rivers (regions). We also used K-H ANOVA to compare C: N: P stoichiometry of cattle dung among regions. Significant H-tests were followed by Tukey multiple comparisons as post hoc tests.

We used generalized additive mixed modelling (GAMM) to test for spatial and seasonal variation in livestock characteristics (number of livestock and percentage of individuals defecating and urinating in the river per herd) using the mgcv package in R (52, 53). Before GAMMs count data were log-transformed while percentage data were logit-transformed. For each response variable, the GAMM model included river (Amala, Nyangores and Talek) and season (dry and wet) as fixed effects, and watering point (sampling site) as a random effect to test whether site location influenced livestock characteristics. We included river and its interaction with the season (river X season) as fixed factors. We fit an initial GAMM ‘full’ model that included river and season as fixed effects, and ‘watering point’ as a random effect. To identify the most parsimonious model we used a step-wise approach based on the Akaike Information Criterion (AIC) to achieve an optimal model (54).

We used bootstrap analysis (k =10,000 with replacement) to estimate 95% confidence intervals (CIs) for livestock characteristics data using the boot package in R (55). Bootstrapping is a resampling method used for estimating a distribution, from which various measures of interest can be calculated (e.g., mean, standard error and CIs) (56–58). We used paired t-tests to compare *in situ* water variables, concentrations of chlorophyll-a, organic matter, dissolved organic carbon and nutrients between upstream and downstream locations at livestock watering points.

## Results

### Cattle behaviour

There were seasonal differences in stream size at watering points brought about by increases in discharge during the wet season (Table 1). The Amala and Nyangores regions had lower numbers of cattle visiting watering points than the Talek region (Figure 3). The median number of cattle per herd in the Nyangores and Amala was 4 and herd size ranged from 1 to 14 in Nyangores and 1 to 18 in Amala, while in the Talek the median was 50, with a range from 4 to 2100. In the Talek, two herds were quite large at 1500 and 2100 individuals and were the only ones with numbers over 600, with the third-highest herd having 530 individuals. There were no significant spatial and seasonal differences in time spent by cattle in the river, and the percentage of cattle per herd that defecated or urinated in the river (Figure 3, Table 2).

**Figure 3.**
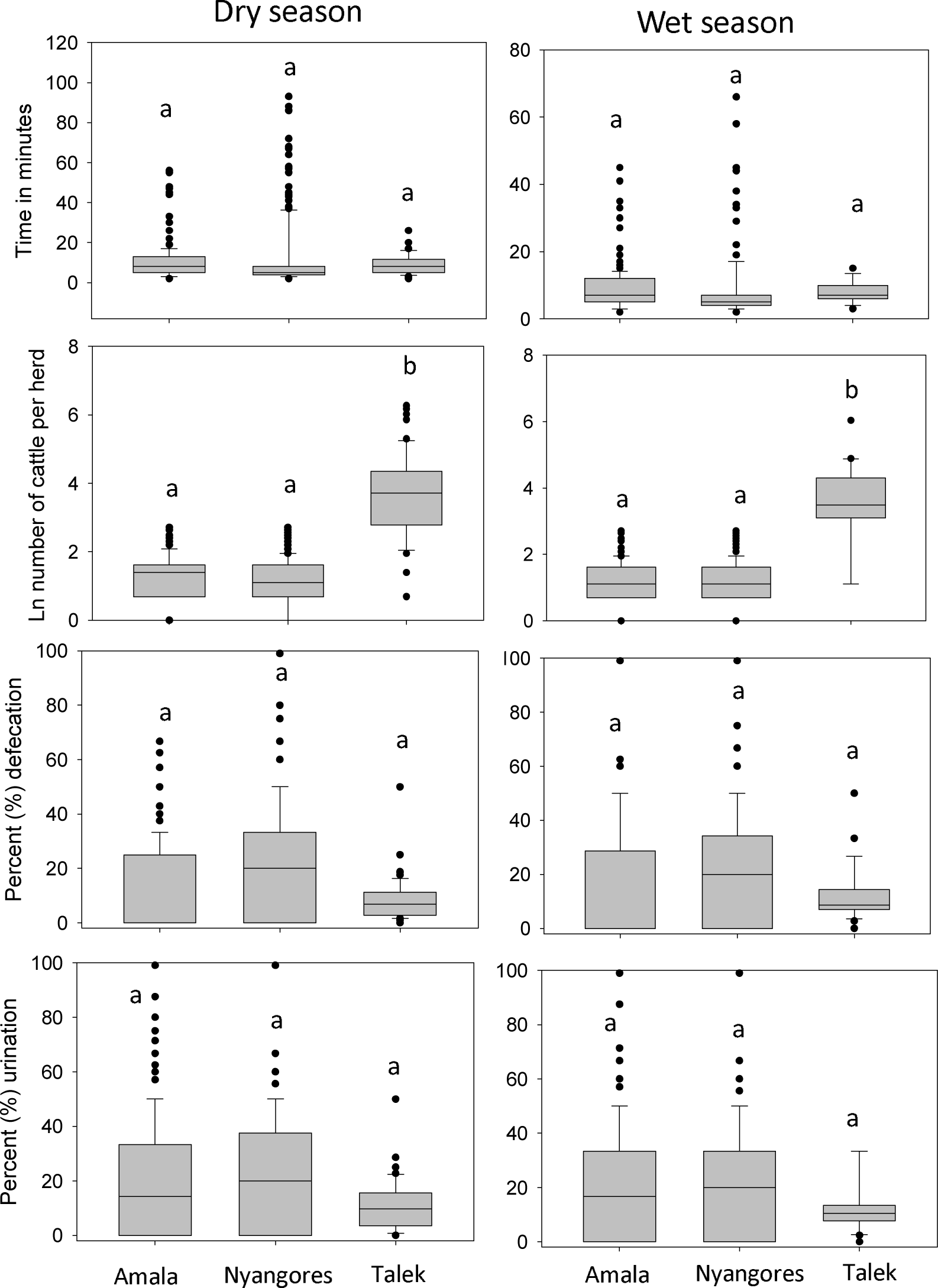
Livestock behaviour (time spent in the watering points, number of cattle per herd, % defecation and % urination) in the upper Mara River Basin and lower basin in the Talek region during the wet and dry seasons.

**Table 1:** Sizes of livestock watering points and total number of cattle that visited watering points in the Mara River Basin, Kenya during the dry and wet seasons. Given are mean value ± standard deviation, H-test after Kruskal-Wallis (***p < 0.001, **p < 0.01, *p < 0.05), except for number of cattle and number of watering points. Lower case superscripted letters indicate significant differences according to Tukey post-hoc tests following a significant H-test.

| Characteristics | Nyangores<br>Dry | Nyangores<br>Wet | Amala Dry | Amala Wet | Talek Dry | Talek Wet | H-test |
| --- | --- | --- | --- | --- | --- | --- | --- |
| Number of watering points | 21 | 16 | 17 | 13 | 10 | 10 | - |
| Total number of cattle | 768 | 558 | 795 | 603 | 7,836 | 1576 | - |
| River width (m) | 3.6±0.9<sup>a</sup> | 4.3±0.6<sup>a</sup> | 3.1±0.7<sup>a</sup> | 4.5±1.2<sup>a</sup> | 4.1±1.0<sup>a</sup> | 9.7±2.4<sup>b</sup> | 0.097 |
| River depth (m) | 0.2±0.1<sup>a</sup> | 0.2±0.1<sup>a</sup> | 0.2±0.05<sup>a</sup> | 0.2±0.1<sup>a</sup> | 0.1±0.03<sup>b</sup> | 0.3±0.1<sup>a</sup> | 0.048* |
| Discharge (m<sup>3</sup>/s) | 0.2±0.1<sup>a</sup> | 0.6±0.2<sup>b</sup> | 0.1 ± 0.1<sup>a</sup> | 0.5 ± 0.3<sup>b</sup> | 0.1±0.6<sup>a</sup> | 1.0±0.6<sup>b</sup> | 0.034* |

**Table 2:** Summary of generalized additive mixed models (GAMMs) used to determine the effect of the river system (region) and seasonality (wet and dry seasons) livestock characteristics: time (minutes) spent at watering points, number of cattle per herd, percentage of cattle defecating or urinating in the river. The ‘full’ model included the river or region (Amala, Nyangores and Talek), season (dry vs. wet) and river X season interaction as fixed effects and watering point as a random effect.

| Fixed effects | Livestock characteristics at watering points |  |  |  |
| --- | --- | --- | --- | --- |
|  | Time spent watering<br>(min) | Number of cattle per<br>herd | Percent defecated | Percent urinated |
| Intercept (estimate (SE); t value) | 1.93(0.31); 6.20*** | 3.88(0.41); 9.56*** | 0.15(0.09); 1.64 | 0.15(0.11); 1.42 |
| River (Estimate (SE); t value) | 0.06(0.13); 0.47 | -1.01(0.17); -5.87*** | -0.01(0.04); -0.16 | 0.02(0.04); 0.49 |
| Season Estimate (SE); t value | -0.11(0.21); -0.54 | -0.06(0.20); -0.31 | 0.02(0.06); 0.34 | 0.004(0.07); 0.06 |
| River x Season (estimate (SE); t value) | 0.02(0.08); 0.22 | 0.03(0.08); 0.33 | 0.01(0.02); 0.24 | 0.001(0.03); 0.02 |
| **Random effect** |  |  |  |  |
| Watering point (intercept) SD | 0.22 | 0.63 | 0.06 | 0.05 |
| Residual SD | 0.72 | 0.70 | 0.21 | 0.26 |
| Adjusted $R^2$ | 0.004 | 0.20 | 0.002 | -0.002 |
| Scale estimation | 0.52 | 0.48 | 0.04 | 0.07 |

SE= standard error; SD = degrees of freedom; t = t-test value between the fitted and a null model. Significance: \* $p < 0.05$ , \*\* $p < 0.01$ , \*\*\* $p < 0.001$

The bootstrap data and 95 confidence intervals (CIs) for livestock characteristics and C, N and P composition of cattle dung and urine are presented in Table S1 (supplementary information). The median time spent by cattle at watering points was 11.5 minutes, and the 95% CIs were 10.6 and 12.5 minutes (Figure 3). Across the MR basin slightly more cattle urinated (bootstrap median = 13.6%) than defecated (bootstrap median = 12.4%) in the river. The bootstrap CIs for defecation and urination were 11.3-13.6% and 12.2-14.9%, respectively. The median dung weight was 868.5 g, and the 95% CIs were 749.8 g and 991.1 g. The median urine volume was 0.788 L, and the 95% CIs were 0.611 L and 0.965 L.

### Loading rates by cattle

There was a significant difference in the C: N: P stoichiometry (quality) of cattle dung among regions (Table 3). The Nyangores and Amala regions recorded the highest quality (lower C: N and C: P ratios) of cattle dung loaded into rivers, while the lower MR basin (Talek and MMNR) recorded the poorest quality (highest C: N ratio). Cattle dung had a lower C: N and C:P ratio (higher quality) than hippo dung (Table 3). On average, the composition of cattle dung was 32.6±4.2% C, 1.4±0.3% N and 0.29±0.07% P, while that of urine was 14.2±2.7 % C, 10.3±1.2% N and 0.43±0.13% P. Cattle dung and urine had a stoichiometry of 113.2 C: 4.9 N: 1.0 P and 33.2 C: 23.9 N: 1.0 P, respectively. Overall stoichiometry by mass of cattle excretion/ egestion in the MR basin was 57.2 C: 19.3 N: 1.0 P.

**Table 3.** Mean (±SD) quality (C: N: P ratio) of cattle dung and urine and hippo dung in different regions in the Mara River basin, Kenya.

| Regions | Cattle |  |  |  | Hippopotamus |  |  |  |
| --- | --- | --- | --- | --- | --- | --- | --- | --- |
|  | C: N | C: P | N: P | C: N: P | C: N | C: P | N: P | C: N: P |
| **Dung** |  |  |  |  |  |  |  |  |
| Nyangores | 18.4±3.9<sup>a</sup> | 97.5±4.4<sup>a</sup> | 5.3±0.6<sup>a</sup> | 97.5:5.3:1 | - | - | - | - |
| Amala | 21.7±4.1<sup>a</sup> | 102.2±5.2<sup>a</sup> | 4.7±0.7<sup>ab</sup> | 102.2:4.7:1 | - | - | - | - |
| Middle Mara | 24.9±4.5<sup>ab</sup> | 117.4±2.3<sup>ab</sup> | 4.7±0.7<sup>ab</sup> | 117.4:4.7:1 | 23.27±3.72<sup>b</sup> | 263.7±8.9<sup>a</sup> | 5.5±0.9<sup>a</sup> | 263.7:5.5:1 |
| MMNR | 26.1±3.7<sup>b</sup> | 127.2±4.2<sup>b</sup> | 4.9±0.5<sup>ab</sup> | 127.2:4.9:1 | 34.27±6.38<sup>a</sup> | 227.5±12.6<sup>a</sup> | 5.9±0.9<sup>a</sup> | 227.5:5.9:1 |
| Talek Region | 27.2±4.9<sup>b</sup> | 122.3±3.1<sup>b</sup> | 4.5±0.4<sup>b</sup> | 122.3:4.5:1 | 33.30±7.58<sup>ab</sup> | 257.4±10.2<sup>a</sup> | 6.6±0.7<sup>a</sup> | 257.4:6.6:1 |
| F - value | 3.69 | 4.34 | 2.68 | - | 2.54 | 1.97 | 1.42 | - |
| P - value | 0.004 | 0.01 | 0.044 | - | 0.041 | 0.088 | 0.231 | - |
| **Urine** |  |  |  |  |  |  |  |  |
| Talek Region | 1.4±0.06 | 36.0±11.0 | 25.7±7.3 | 33.2:23.9:1 | - | - | - | - |

Based on metabolic considerations, we estimate that cattle in the MR basin had a daily intake of 25 g dry matter (DM) kg^-1^ in the dry season and 19 g DM kg ^-1^ in the wet season. This translates to an egestion of 10.5 g DM kg cattle^-1^ day^-1^ in the wet season, and 13.8 g DM kg cattle^-1^ day^-1^ in the dry season. Assuming that cattle consumption is averaged over 6 months of the wet season and 6 months of the dry season and that they spend 11.5 minutes in the river per day, we estimated that individual cattle with a body mass of 265 kg loads 25.6 g DM kg day^-1^ into the river. With a wet-dry mass conversion of 25.7%, we estimated that per head, 99.6 g (wet mass) of OM (dung) is loaded into rivers per day. Per capita, cattle defecate 12.5 kg faeces (wet mass) every day, and 0.0996 kg (0.7%) of that goes into the Mara River. Assuming an average daily urine volume of 6.63 L and a time of 11.5 minutes spent in or near the river, we estimate a daily per capita urine volume deposited in or near the river of 0.053 L.

Direct observations of defecation yielded an average of 868.5±7.9 g wet mass (223.2 g DM) of dung per defecation event, and this enters the river every time cattle defecates in or near the river. Since out of a herd of cattle that visit a watering point, on average only 12.4% defecate, this translates to per capita faeces (wet mass) deposition in or near rivers of 107.7 g, which is marginally higher than the estimate of 99.6 g faeces based on metabolic considerations. Further, 13.6% of the herd were observed to urinate during the visits to watering points. Assuming a volume of 6.63 L per single urination event, this translates to a per capita urine loading of 0.090 L. This estimate is based on the number of observed urination events (10) per day and their average volume, which is 70% higher than the estimate based on fractional time spent near the watering point and the daily urine production.

We used only the metabolism-based loading estimates for further computation of C, N and P fluxes upscaled to region-wide estimates. The motivation behind this choice is for better comparability with the previously published hippo-driven fluxes, which were achieved using the same methodology, and ease of gaining further data in future projects.

Based on metabolism model, per capita cattle added 99.6 g wet mass (25.6 g DM) of OM, 8.4 g C, 0.4 g N and 0.07 g P through egestion, and 7.6 g C, 5.5 g N and 0.23 g P through excretion into the Mara River per day (Table 4). The overall loading of waste (excretion + egestion) per cattle per day into the Mara River was 99.6 g OM (wet mass), 16.0 g C, 5.9 g N, and 0.3 g P per day (Table 4).

**Table 4.** Per capita loading rates for organic matter (OM), carbon (C), nitrogen (N) and phosphorus (P) through egestion and excretion by cattle determined by the metabolism model and direct method for the Mara River basin, Kenya. 95% confidence intervals for cattle dung loading rates are provided in brackets.

| Loading rates by cattle | Method of estimation |  |
| --- | --- | --- |
|  | Metabolism model | Direct method |
| **Cattle dung** |  |  |
| OM (wet mass, (g cattle<sup>-1</sup> day<sup>-1</sup>)) | 99.6 (91.8-108.3) | 107.7 (84.8-134.8) |
| C (g cattle<sup>-1</sup> day<sup>-1</sup>) | 8.4 (7.7-9.1) | 9.02 (6.6-12.1) |
| N (g cattle<sup>-1</sup> day<sup>-1</sup>) | 0.4 (0.3-0.4) | 0.4 (0.4-0.6) |
| P (g cattle<sup>-1</sup> day<sup>-1</sup>) | 0.07 (0.07-0.09) | 0.08 (0.07-0.11) |
| **Cattle urine** |  |  |
| C (g cattle<sup>-1</sup> day<sup>-1</sup>) | 7.6 (6.6-9.6) | 12.8 (8.7-18.3) |
| N (g cattle<sup>-1</sup> day<sup>-1</sup>) | 5.5 (4.7-7.1) | 9.2 (6.3-13.5) |
| P (g cattle<sup>-1</sup> day<sup>-1</sup>) | 0.23 (0.16-0.31) | 0.39 (0.21-0.66) |
| **Total egestion and excretion** |  |  |
| OM (wet mass, (g cattle<sup>-1</sup> day<sup>-1</sup>)) | 99.6 | 107.7 |
| C (g cattle<sup>-1</sup> day<sup>-1</sup>) | 16.0 | 21.8 |
| N (g cattle<sup>-1</sup> day<sup>-1</sup>) | 5.9 | 9.6 |
| P (g cattle<sup>-1</sup> day<sup>-1</sup>) | 0.3 | 0.5 |

### Cattle as a replacement for wildlife

Using the metabolism method, specific C: N: P stoichiometry of cattle and hippo dung per region and cattle population numbers, we estimated total daily loading of organic matter (OM) for the cattle population in the MMNR to be 1,157 kg. In the Middle Mara and Talek regions, the loading rates for OM were estimated to be 2,599 and 7,364 kg faeces (wet mass), respectively (Table 5). In comparison, total daily loading of OM by the hippopotamus population in the MMNR, Middle Mara and Talek Region, is estimated to be 16,739 kg, 13,668 kg and 5,638 kg faeces (wet mass), respectively (4). Thus, within the MMNR, livestock loading with OM is only 7% of loading originating from hippos, but in the Middle Mara and Talek Region, the loading by cattle increases to 19% and 131%, respectively. These numbers describing the effects of replacing wildlife by livestock change markedly when the differences of cattle vs. hippos concerning wet-dry conversion factors for dung and stoichiometry of egestion and excretion are taken into account. Daily loading of C, N and P due to the cattle in the MMNR represents 12.1% C, 29.5% N and 15.8% P of the loading achieved by hippopotamus (Table 5). These relative loading rates for cattle vs. hippos increase to 31.8% C, 80.9% N and 43.6% P in the Middle Mara region, and to 224.6% C, 556.7% N and 274.4% P in the Talek region. Overall, regarding OM loading, one hippo corresponds to the loading of 100 individuals of cattle, while for C, N and P loading it corresponds to an equivalent of 59, 24 and 44 cattle, respectively.

**Table 5.** Median loading rates of organic matter (dung) and nutrients (C, N and P) by cattle in the Mara River basin based on the metabolism model in comparison with published loading rates for hippopotamus. Cattle numbers outside the reserve area for the Koyake Group Ranch, while numbers for the Talek River represent all other Group Ranches, estimated from a conservative number of 100,000 cattle in the group ranches outside the MMNR. ^y^Published hippopotamus loading rates are from (4).

| **Cattle and hippo populations and loading numbers** | **Nyangores** | **Amala** | **Middle Mara** | **Talek Region** | **MMNR** |
| --- | --- | --- | --- | --- | --- |
| Cattle numbers | 16285 | 21581 | 30,000 | 85,000 | 13,350 |
| **Cattle dung** |  |  |  |  |  |
| Loading by cattle population (OM wet mass kg day<sup>-1</sup>) | 1410.9 | 1869.8 | 2599.2 | 7364.4 | 1156.6 |
| Loading by cattle population (C in kg day<sup>-1</sup>) | 105.7 | 138.5 | 224.5 | 669.1 | 109.2 |
| Loading by cattle population (N in kg day<sup>-1</sup>) | 7.3 | 6.3 | 8.8 | 24.9 | 3.8 |
| Loading by cattle population (P in kg day<sup>-1</sup>) | 1.1 | 1.5 | 1.9 | 5.1 | 0.8 |
| **Cattle Urine** |  |  |  |  |  |
| Loading by cattle population (C in kg day<sup>-1</sup>) | 107.1 | 142.0 | 197.4 | 559.2 | 87.8 |
| Loading by cattle population (N in kg day<sup>-1</sup>) | 77.3 | 102.5 | 142.5 | 403.7 | 63.4 |
| Loading by cattle population (P in kg day<sup>-1</sup>) | 3.2 | 4.3 | 6.0 | 16.9 | 2.6 |
| **Total egestion and excretion** |  |  |  |  |  |
| Total loading by cattle population OM (wet mass, (g cattle<sup>-1</sup> day<sup>-1</sup>) | 1410.9 | 1869.8 | 2599.2 | 7364.4 | 1156.6 |
| Total loading by cattle population C (g cattle<sup>-1</sup> day<sup>-1</sup>) | 212.9 | 280.5 | 421.8 | 1228.3 | 197.0 |
| Total loading by cattle population N (g cattle<sup>-1</sup> day<sup>-1</sup>) | 84.7 | 108.8 | 151.3 | 428.6 | 67.2 |
| Total loading by cattle population P (g cattle<sup>-1</sup> day<sup>-1</sup>) | 4.4 | 5.8 | 7.9 | 22.0 | 3.5 |
| **Total loading by hippopotamus population** |  |  |  |  |  |
| Hippopotamus numbers | - | - | 1,571 | 648 | 1,924 |
| OM (wet mass in kg day<sup>-1</sup>)<sup>y</sup> | - | - | 13,668 | 5,638 | 16,739 |
| C (g hippopotamus<sup>-1</sup> day<sup>-1</sup>) | - | - | 1,327 | 547 | 1,625 |
| N (g hippopotamus<sup>-1</sup> day<sup>-1</sup>) | - | - | 187 | 77 | 228 |
| P (g hippopotamus<sup>-1</sup> day<sup>-1</sup>) | - | - | 18 | 8 | 22 |

Loading rates for OM and nutrients were also estimated per unit area of the river. Using the average widths of the Mara River and its major tributary the Talek River in its lower section (20 and 10 m, respectively), on average cattle load a total of 457.5 g DM, 75.8 g C, 26.6 g N and 1.4 g P m^-2^ year^-1^ into riverine habitats of the lower MR basin (Mara River, lower Talek and Olare-Orok tributaries). Along the upper Talek River where livestock densities are very high, cattle loading increases by >100% to 1193.6 g DM, 199.1 g C, 69.5 g N, and 3.6 g P m^-2^ year.

### Livestock effects on water quality

Downstream locations of livestock watering points recorded significantly higher TSS and POM concentrations compared to upstream locations (paired t-test, p <0.05, Table 6). However, no significant differences were noted for temperature, DO, EC, TDS, pH, water column chlorophyll-a and benthic chlorophyll-a. For nutrients in the water column, mean concentrations were higher downstream, but only significantly for TP, TN and TDN, with relevant increases of 54% and 44% for N fractions, respectively. Differences between upstream and downstream locations were more pronounced for nutrients in the sediments, with nitrate showing the highest increase of 60% (Table 6). To capture the direct effects of livestock presence on water quality, differences in diel (morning, nooon/mid-day and evening) levels of physico-chemical variables and concentrations of nutrients were used (Figure 4 and 5). As expected, we reccorded higher mean water temperature and lower dissolved oxygen concentrations, but no differences in electrical conductivity, total dissolved soilds and salinity (Figure 4). However, there were clear diel changes in nutrient concentrations occassioned by the presence of livestock at the watering points (Figure 5). For instance, nitrates, ammonia and soluble reactive phosphorus concentrations were higher during mid-day than the rest of the times (morning and evening), and for most nutrients, dowstream concentrations were higher than upstream concentrations (Figure 5).

**Figure 4.**
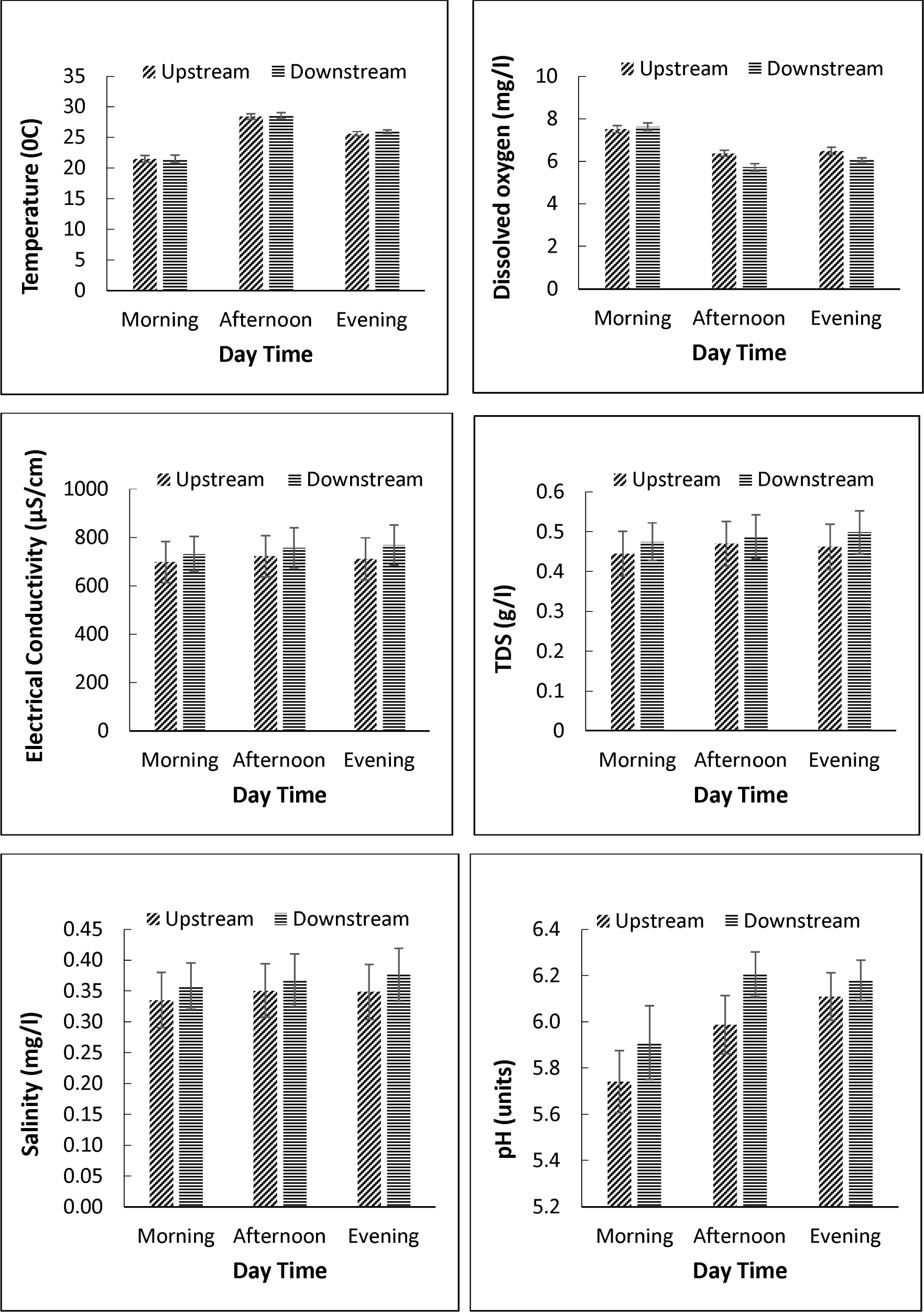
Changes in *in-situ* physico-chemical parameters during different times of the day at livestock watering points in the Mara River basin, Kenya. The different times correspond to diel (morning, noon/mid-day and evening) variation in the number of livestock visiting watering points.

**Figure 5.**
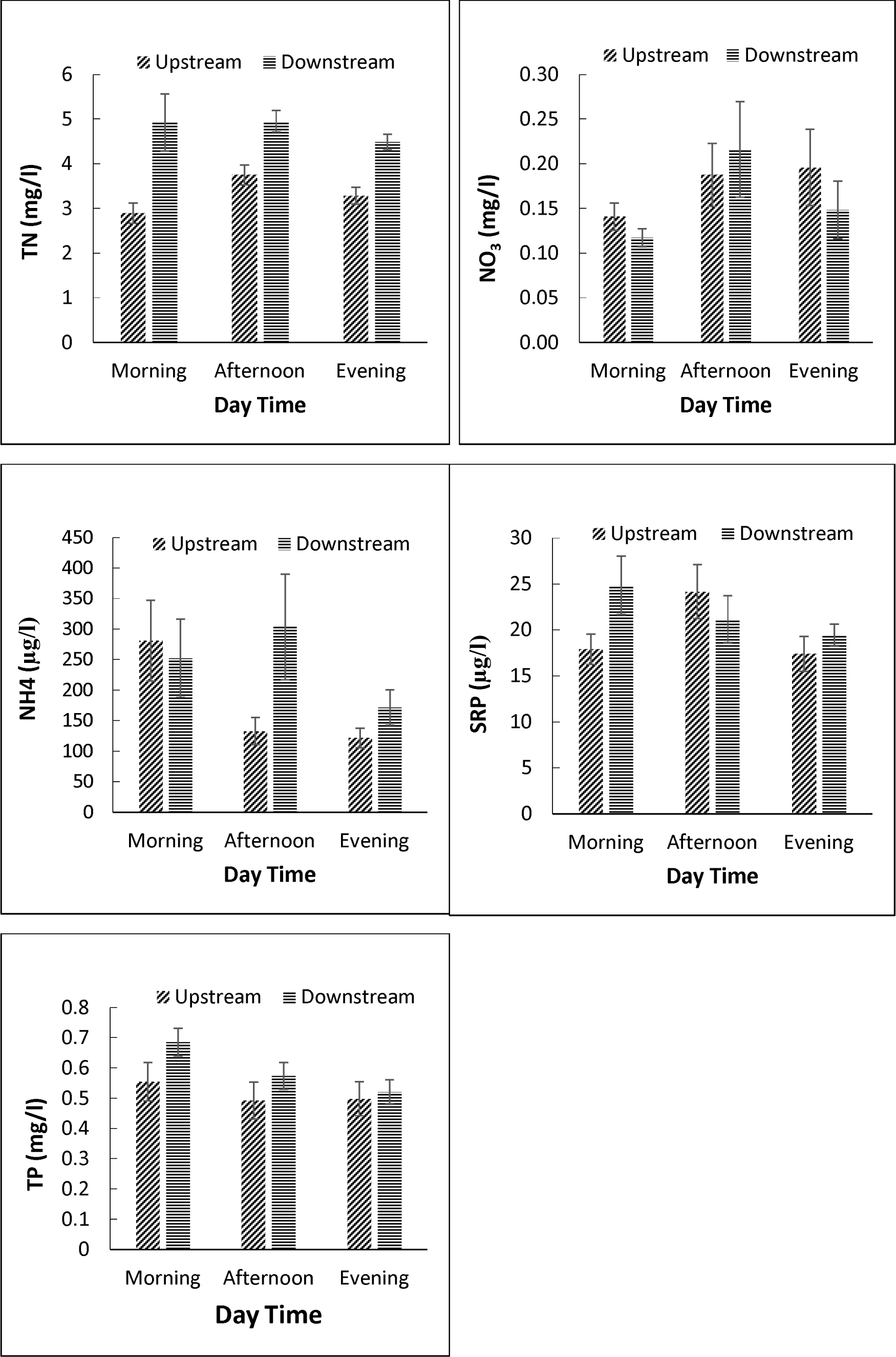
Changes in nutrient concentrations during different times of the day at livestock watering points in the Mara River basin, Kenya. The different times correspond to diel (morning, noon/mid-day and evening) variation in the number of livestock visiting watering points.

**Table 6:**
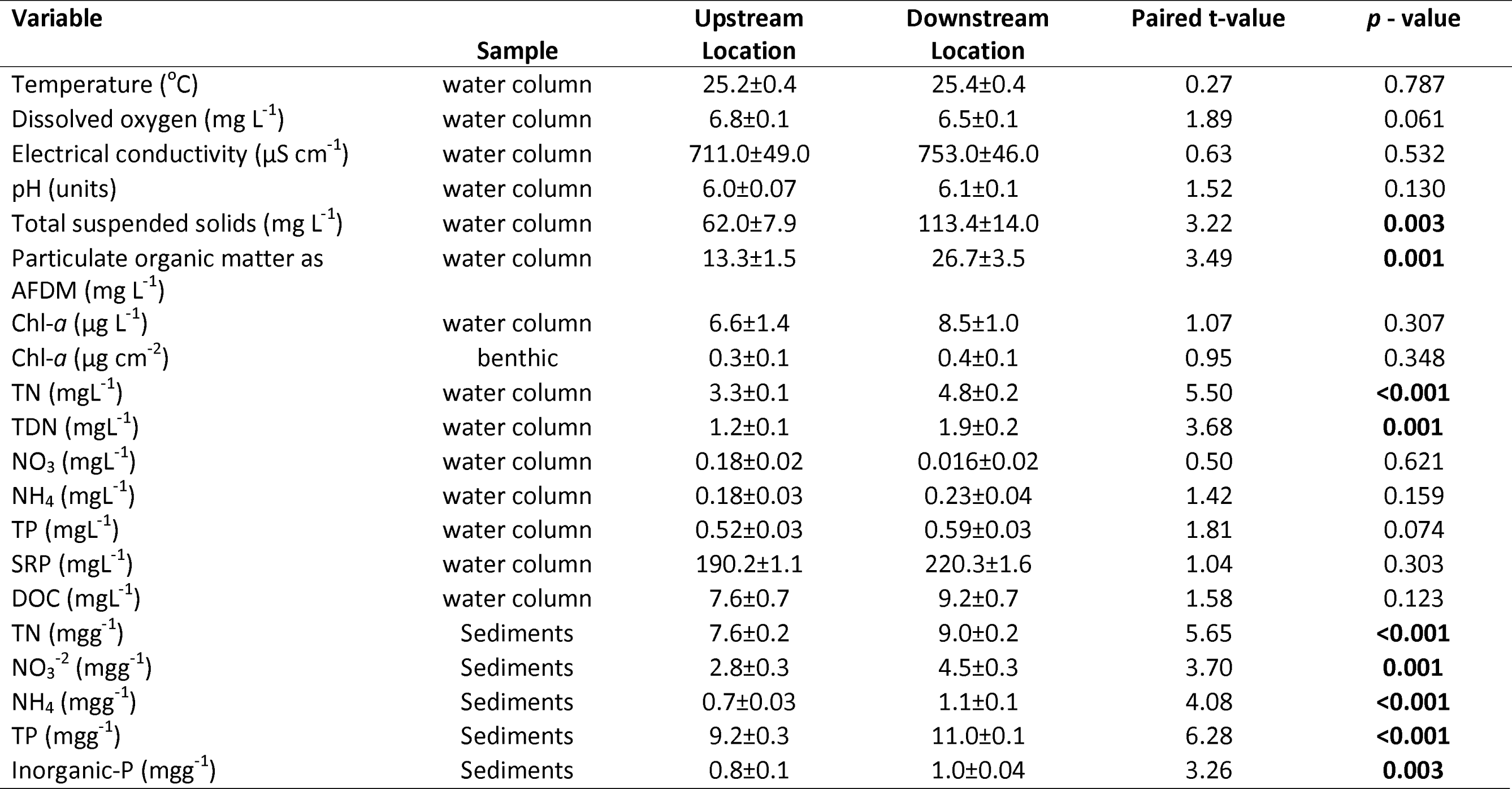
Mean (±SD) variation in water quality variables and nutrient concentrations in the water column and sediments at the upstream and downstream river reaches of livestock watering sites in the Talek River, a tributary of the Mara River.

| Variable | Sample | Upstream Location | Downstream Location | Paired t-value | *p* - value |
| --- | --- | --- | --- | --- | --- |
| Temperature ($^{\circ}$C) | water column | 25.2±0.4 | 25.4±0.4 | 0.27 | 0.787 |
| Dissolved oxygen ($\text{mg L}^{-1}$) | water column | 6.8±0.1 | 6.5±0.1 | 1.89 | 0.061 |
| Electrical conductivity ($\mu\text{S cm}^{-1}$) | water column | 711.0±49.0 | 753.0±46.0 | 0.63 | 0.532 |
| pH (units) | water column | 6.0±0.07 | 6.1±0.1 | 1.52 | 0.130 |
| Total suspended solids ($\text{mg L}^{-1}$) | water column | 62.0±7.9 | 113.4±14.0 | 3.22 | **0.003** |
| Particulate organic matter as AFDM ($\text{mg L}^{-1}$) | water column | 13.3±1.5 | 26.7±3.5 | 3.49 | **0.001** |
| Chl-*a* ($\mu\text{g L}^{-1}$) | water column | 6.6±1.4 | 8.5±1.0 | 1.07 | 0.307 |
| Chl-*a* ($\mu\text{g cm}^{-2}$) | benthic | 0.3±0.1 | 0.4±0.1 | 0.95 | 0.348 |
| TN ($\text{mg L}^{-1}$) | water column | 3.3±0.1 | 4.8±0.2 | 5.50 | **<0.001** |
| TDN ($\text{mg L}^{-1}$) | water column | 1.2±0.1 | 1.9±0.2 | 3.68 | **0.001** |
| NO<sub>3</sub> ($\text{mg L}^{-1}$) | water column | 0.18±0.02 | 0.016±0.02 | 0.50 | 0.621 |
| NH<sub>4</sub> ($\text{mg L}^{-1}$) | water column | 0.18±0.03 | 0.23±0.04 | 1.42 | 0.159 |
| TP ($\text{mg L}^{-1}$) | water column | 0.52±0.03 | 0.59±0.03 | 1.81 | 0.074 |
| SRP ($\text{mg L}^{-1}$) | water column | 190.2±1.1 | 220.3±1.6 | 1.04 | 0.303 |
| DOC ($\text{mg L}^{-1}$) | water column | 7.6±0.7 | 9.2±0.7 | 1.58 | 0.123 |
| TN ($\text{mg g}^{-1}$) | Sediments | 7.6±0.2 | 9.0±0.2 | 5.65 | **<0.001** |
| NO<sub>3</sub><sup>-2</sup> ($\text{mg g}^{-1}$) | Sediments | 2.8±0.3 | 4.5±0.3 | 3.70 | **0.001** |
| NH<sub>4</sub> ($\text{mg g}^{-1}$) | Sediments | 0.7±0.03 | 1.1±0.1 | 4.08 | **<0.001** |
| TP ($\text{mg g}^{-1}$) | Sediments | 9.2±0.3 | 11.0±0.1 | 6.28 | **<0.001** |
| Inorganic-P ($\text{mg g}^{-1}$) | Sediments | 0.8±0.1 | 1.0±0.04 | 3.26 | **0.003** |

## Discussion

### Loading of livestock and hippos

Our findings show that cattle are major agents for the transfer of organic matter, carbon and nutrients (N and P) from terrestrial to aquatic environments in African savannas and grazing areas. On average, an individual zebu cattle contributes 36.4 kg of OM (wet weight) in the form of dung, 5.8 kg C, 2.2 kg N and 0.11 kg P year^-1^ into the Mara River through excretion and egestion. Given the 1.8 million cattle population in the MR basin, we estimate total daily loading into the river to be 179.7 metric tons OM (wet mass), 28.3 metric tons C, 10.5 metric tons N and 0.6 metric tons P. In comparison, daily loading by the hippopotamus population (approximately 4000 individuals) into the river is approximately 36.2 metric tons OM (wet mass), 3.5 metric tons C, 0.5 metric tons N and 0.05 metric tons P. Cattle inputs of OM are estimated to range from 7% -131% of loading relative to hippopotamus loading rates in areas where their distribution overlap.

Arguably, these numbers are first-order estimates as they rely on several assumptions. For example, a daily visit of hippo to a watercourse is guaranteed but may be doubted for cattle. Also, it is estimated that livestock watering in the river occurs only once during the day, but cattle are sometimes watered or cross the river twice when leaving for grazing and returning to bomas (livestock sheds or enclosures) in the evening. There is significant spatial and temporal variation in loading rates as a result of spatial variations in cattle densities, forage availability and quality and distribution of water sources and distance covered or time spent foraging. Other factors that influence cattle loading to the river include grassland productivity, which is highly dependent on seasonal and annual variations in precipitation and grazing intensity (59).

There was spatial variation in the C: N: P stoichiometry of dung across the MR basin with the upper basin (Nyangores and Amala) having lower C relative to N and P than the lower basin (Talek region and MMNR) (Table 3). While this may be indicative of changes in forage composition among regions, it also could be due to differences in grazing regimes. It is notable that in the agricultural areas (Nyangores, Amala and Upper Mara) where mixed crop and livestock farming is practised, livestock feed on other types of forage other than pasture (grass), such as Napier grass and maize stalks. In comparison, livestock in the Middle Mara, Talek Region and MMNR mainly forage on savanna grass with limited access to supplementary feeds. The carbon content of dung can vary strongly due to variation in organic matter content, feed digestibility and feed quality (C: N: P ratio), thereby also affecting N and P content. Also, while the small paddocks in the upper MR basin are intensively grazed and pasture is dominated by fresh shoots of grasses, the middle Mara, MMNR and Talek regions are mainly composed of tussock that is of poorer quality. Low C: N ratios in dung, hence in forage (grass) in the upper MR basin could result from accelerated nutrient cycling or increased nutrient availability induced by livestock faeces and urine (60)

Variation in ration digestibility (quality) and protein content can also result in large variations in nitrogen excretion and egestion (61, 62). The C: N stoichiometry of dung (range 18.4±3.9 -27.2±4.9) obtained in this study are within ranges reported for African cattle or some grazers at pasture (63, 64). Low C: N ratio in dung is indicative of high-quality forage or feeds that are rich in protein, while high C: N and C: P ratios are indicative of low-quality forage that is typical of savanna grass during the dry season (10). Similar findings of a high C: N and C: P ratios have also been reported for cattle dung in semi-arid eastern Kenya (65). Although we did not consider seasonality in our study, the composition of dung and urine can vary substantially between seasons due to differences in feed availability and quality (66). However, some studies have reported limited or lack of variation in C: N ratio of dung between seasons in savanna grasslands in Zimbabwe (10).

In our study, cattle spent an average of 11.5 minutes, which is 2% of the total observation time (9:00-18:00 hrs), in or near the river during watering or crossings. In a similar study, Bond et al. (23) observed that cattle spend approximately 2% and 7% of their time (8:30-16:00 hrs) in the aquatic environment and riparian zone, respectively. Other studies have reported less time spent by cattle in or near aquatic environments. Ballard and Krueger (67) recorded 1%, whilst Haan et al. (68) recorded the duration of in-stream cattle activity to be 1.1%. These differences can be explained by factors such as methodological aspects, herding and environmental differences among studies. Methodologically, both Ballard and Krueger (67) and Haan et al. (68) used an insufficiently frequent recording interval for observation, while Bond et al. (2014) used continuous observation as we did in this study. Also, while cattle were left to roam freely and visit the river or watering points without restrictions, most of the river visits by cattle in our study are largely decided by herders.

Because of differences in cattle stocking densities and C: N: P stoichiometry of dung, the average areal loading rates we estimated for cattle differed between the upper MR basin (229.5 g DM, 35.1 g C, 13.5 g N and 0.7 g P m^-2^ year^-1^) and the lower MR basin (702.9 g DM, 116.9 g C, 40.9 g N and 2.1 g P m^-2^ year^-1^). In the MR basin, the distribution of cattle is not uniform and some regions, such as the upper MR basin in Nyangores and Amala where farmers practice mixed crop farming as well as animal husbandry, loading rates are much lower than to the lower MR basin, where the Maasai pastoralists keep large numbers of cattle. For both regions, these estimates are lower than estimates for some wild LMH in African savannas. In the Mara River, it has been estimated that hippopotamus loading amounts to 1229 g DM, 502 g C, 71 g N and 6.9 g P m^-2^ year^-1^ (4). In the eulittoral zone of a waterhole in Hwange National Park, Zimbabwe loading by LMH was estimated to be 3157 C, 91 N, and 22 P g m^-2^ year^-1^ (10). On the other hand, cattle loading values in our study are higher than estimates in an English Chalk stream, where loading rates through defecation by 33 cattle was 198 g DM, 8.0 g N and 14.7 g P m^-2^ year^-1^ (9). Differences in loading rates among animal vectors are largely due to differences in body size, whereby megaherbivores such as elephants and hippopotamus consume (and transfer) large amounts of terrigenous vegetation. Some animals spend much more time in and around the river than others, such as hippopotamus, and this increases the loading of organic matter and nutrients. Loading rates are also a function of animal populations and their distribution in river networks.

Our loading estimates for faeces based on metabolic considerations and direct method considerably agree, as opposed to estimates for urine (Table 4). The discrepancies in urine are probably due to relying on literature to estimate urination events per day. Data on frequencies of urination among African zebu cattle are limited, but the value we used for daily urinary output (6.63 L per day) for African zebu cattle agrees with zebu Tharparkar cattle (6.9 L per day) (69), but slightly lower than Nellore zebu cattle (8.1 L per day) (70). It has also been noted that cattle preferentially defecate and urinate in aquatic environments (23), which implies that the volume per urination is likely lower during watering than during other times because the motivation is the trickling stream rather than a full bladder. This agreement in urinary outputs implies that the frequency of urination and diel variation in urination volumes probably presents sources of uncertainty in our study. In a review, Selbie et al. (71) noted that daily urine volume varies widely among different types of cattle, with average volume per urination event ranging from 0.9 L to 20.5 L. Although we estimated ten (10) urination events in our study, with a range of 8-12 urinations per day (72), these estimates are from studies conducted in the temperate zone where water intake and average weights of cattle are likely higher. In semi-arid African savannas, water is limited during the dry season and this will reduce the intake and excretion rates, including the frequency of urination.

### Livestock influences on water quality

There were significant differences in nutrient concentrations in the water column and sediments between upstream and downstream of livestock watering points (Table 6). Elevated concentrations of TSS and POM were also recorded downstream of livestock watering points in the water column. In a similar study, Bond et al. (9) showed that cattle access to a river led to instream increase of nitrogen, phosphorus and potassium concentrations. Through their instream activity, livestock contributes to organic matter and nutrient input and re-suspension of sediments, therefore elevating turbidity levels in streams and rivers (13). In many studies, sediment losses from trampled and heavily grazed stream banks have been reported to exceed those observed for untrampled or ungrazed counterparts (73).

Well researched aspects of livestock effects on aquatic ecosystems have done in large rangelands North America (e.g., (12, 74) and dairy farms in Australia (e.g., (75) and New Zealand (e.g. (24) where stock densities, management practices and climatic conditions are different from African savannas. Moreover, a wide body of research has shown that cattle access to streams and rivers can have potentially harmful effects on aquatic ecology, geomorphology and water quality. Herding of livestock near the river channels can cause bank slumping or collapse, releasing significant amounts of sediments (76, 77). These livestock-induced habitat changes degrade water quality, alter instream habitats and reduce biodiversity of macroinvertebrates and fishes (13, 78, 79). The most noted effects of stream degradation caused by livestock activity have been the elimination of sensitive macroinvertebrate taxa (e.g., Ephemeroptera, Plecoptera and Trichoptera) and the increase of tolerant species (e.g., Oligochaeta and Chironomidae) (74, 80, 81). Similar effects of hippopotamus on reduced diversity of macroinvertebrates and fishes in streams and rivers have also been reported (16, 82).

Previous studies in the MR basin have judged sediments and nutrient inputs into streams and rivers from livestock grazing and agricultural land use as non-point source pollution (83–85), but other studies have underlined the immediate, local effects of wildlife (hippopotamus) on suspended sediments, organic matter and nutrient input (15, 18, 86). We found notable differences in N and P concentrations in the water column and benthic sediments between the upstream and downstream reaches, with remarkably higher concentrations in the sediments downstream. This shows that cattle can produce similarly localized changes in water quality as hippos and thus, contribute to heterogeneity in the aquatic ecosystem in a similar way as achieved by hippo pools with distinct locations in the landscape. Watering points in our study were rarely locations of specifically high residence time that would smooth and potentially amplify a subsidy effect, but still, differences in water chemistry were noted during different times of the day as a result of direct livestock activity in the rivers (Figures 4 and 5). Moreover, sediments downstream of livestock watering points were indicative of a subsidy, likely through their ability to adsorb inputs of P on mineral surfaces and N and P in microbial biomass. These findings are reflective of the catchment-scale effects of livestock grazing and hippo populations on water quality in the Mara River and its tributaries. Historical data from different sites have shown that high livestock and hippo densities are associated with elevated levels of nutrients, dissolved organic carbon and electrical conductivity (Table S2). These findings fit into previous published results on the influence of livestock and other farming practices (including mixed crop framing and livestock rearing) on water quality in the Mara River basin (81, 83–85). In this study, we note that water quality effects were revealed even for relatively minor watering points attracting 40-100 cattle daily.

### Broad implications of this study

This research increases our knowledge about the amount of resource subsidies cattle can transfer from terrestrial into aquatic ecosystems in savanna landscapes, and how these amounts compare with those of large wildlife. As elsewhere, the remaining populations of wild LMH in African savannas and grasslands are only a fraction of the large numbers that were once key features of these landscapes but have been decimated by human settlements, agricultural activities and replacement by livestock (32, 33). Given the critical role that wild LMH have played for millennia connecting aquatic ecosystems with their terrestrial surroundings, there is concern that this important ecological role may be lost. Alternatively, livestock may provide a functional replacement for LMH, thereby maintaining large-scale ecological mechanisms crucial for savannas. Quantitatively, our study shows that a one-on-one replacement of loading by wild LMH, especially hippopotamus, by cattle is unlikely. Per capita loading of OM, C, N and P by small-bodied Zebu cattle is much lower than loading by the mega hippopotamus. However, our results show that larger cattle populations can create substantial terrestrial-aquatic subsidy fluxes. Even with increasing numbers of livestock, the nature of their distribution and behaviour implies that considerable livestock management would be needed to achieve effects per unit area that are comparable to replaced wildlife. For instance, cattle visit watering points only for a short period (11 minutes) compared to 12 h for hippopotamus. Cattle are also distributed throughout the entire area, distributing faeces and urine over a large area in comparison with hippopotamus that resides in groups in specific pools. Nevertheless, herders determine movements and interactions with water sources and livestock management recommendations could be guided along with ideas of ecological replacement keeping alive the functioning of a landscape, rather than guidance by the simplified objective of maintaining unnatural good water quality.

A particular challenge of such efforts exists in the many differences in physiology, foraging behaviour and body sizes between livestock and wildlife, which have a strong bearing on the composition of their material subsidies, and consequently their influence on ecosystem processes. Ruminants such as cattle, goats and sheep have a relatively efficient digestive system compared to non-ruminants such as hippos and elephants, and this difference in digestion produces smaller fecal particle sizes in ruminants (87). However, non-ruminants have longer mean retention times than ruminants, which enhances nutrient extraction from ingesta compared to ruminants. For instance, the overall C: N: P stoichiometry of cattle dung in this study is 113.8: 4.9: 1.0, while that of the hippopotamus is 249.8: 5.9: 1.0. With most of the large wildlife being replaced by ruminants such as cattle, sheep and goats (20, 33, 88), there is potential for a shift in the functioning of aquatic ecosystems as a result of changes in the quality and quantity of subsidies they are receiving. For instance, the differences in the quality of inputs between livestock (mainly ruminants) and hippopotamus (non-ruminants) have been found to produce differences in ecosystem responses, with cattle transferring higher amounts of limiting nutrients (N and P), major ions, and dissolved organic carbon to aquatic ecosystems relative to hippopotamus (22). The higher quality (lower C: N: P ratio) of cattle vs hippo dung has been observed to promote higher primary production in both the benthos and in the water column (22). On the other hand, the larger particles of hippo dung tend to sink to the bottom of aquatic ecosystems where they smoother and reduce benthic production (15, 18, 89). Thus, replacement of wildlife (hippopotamus) by livestock (mainly cattle) will likely stimulate more algal production than the heterotrophic component (bacteria/fungi), and hence shift aquatic ecosystem towards autotrophy.

### Conclusions

With large wildlife in decline and livestock (cattle, goats and sheep) numbers increasing in the African savannas and elsewhere, the quantity and quality of dung being produced will increasingly become a determinant factor on ecosystem productivity and function. Cattle and hippopotamus differ in the amount of organic matter and nutrients they transfer into the aquatic environment. Cattle dung and hippo dung also differ in where they are initially deposited on the landscape, with 50% of hippo dung often deposited directly into the river or on the riverbank, while only 0.7% of cattle dung is deposited in the river. Changing these patterns of organic matter transport and cycling will have significant effects on the structure and functioning of both terrestrial and aquatic ecosystems.

### Data Availability

The data underlying this paper have been submitted the data to the Dryad repository with the following DOI: https://doi.org/10.5061/dryad.d2547d82z.

## Acknowledgements

We are grateful to Gilbert Geemi, Decla Chemutai and Evans Ole Keshe for assistance during field data collection. We are grateful to Henry Lubanga and Augustine Sitati for assistance with sample analysis at the University of Eldoret. We acknowledge Sylvia Ortmann (Leibniz Institute for Zoo and Wildlife Research) for additional chemical analysis of hippo and cattle dung samples. This work was supported by the International Foundation for Science (Research Grant No. A/5810-1); an Alexander von Humboldt Postdoc fellowship to FOM; and an Austrian Development Cooperation fellowship for MSc to JOI.

## Author contributions

JI designed the study, collected field data, performed data analysis and drafted the manuscript. TH designed the study and critically revised the manuscript; GS designed the study, performed data analysis and critically revised the manuscript; FM conceived the study, collected field data, performed data analysis and critically revised the manuscript. All authors gave final approval for publication and agree to be held accountable for the work performed therein.

## Figure Legends

S1 Figure. Time series of discharge data for the major tributaries of the Mara River, Kenya, at the 1LB02 Gauging Station on the Nyangores River (upper panel) and at the 1LA03 Gauging Station on Amala River (lower panel). The red horizontal line on the figures indicated the study period.

S1 Table: Bootstrap medians and 25% and 95% confidence intervals for cattle characteristics and proportions of C, N and P in dung and urine in the Mara River basin, Kenya.

S2 Table. Differences in density of cattle and hippopotamus, and physico-chemical characteristics (mean ± SD) across different sites in the Mara River, Kenya, grouped into five categories: Forested, Agricultural, low density (LD) livestock (mainly cattle), high density (HD) livestock, and hippopotamus (hippos). The statistics are for one-way ANOVA used to analyse significant differences in physical and chemical variables and nutrient concentrations across the five site categories in the Mara River basin, Kenya.

