## Supplementary figures and images for "Livestock as vectors of organic matter and nutrient loading in aquatic ecosystems in African savannas"

### Supplementary Figure 1

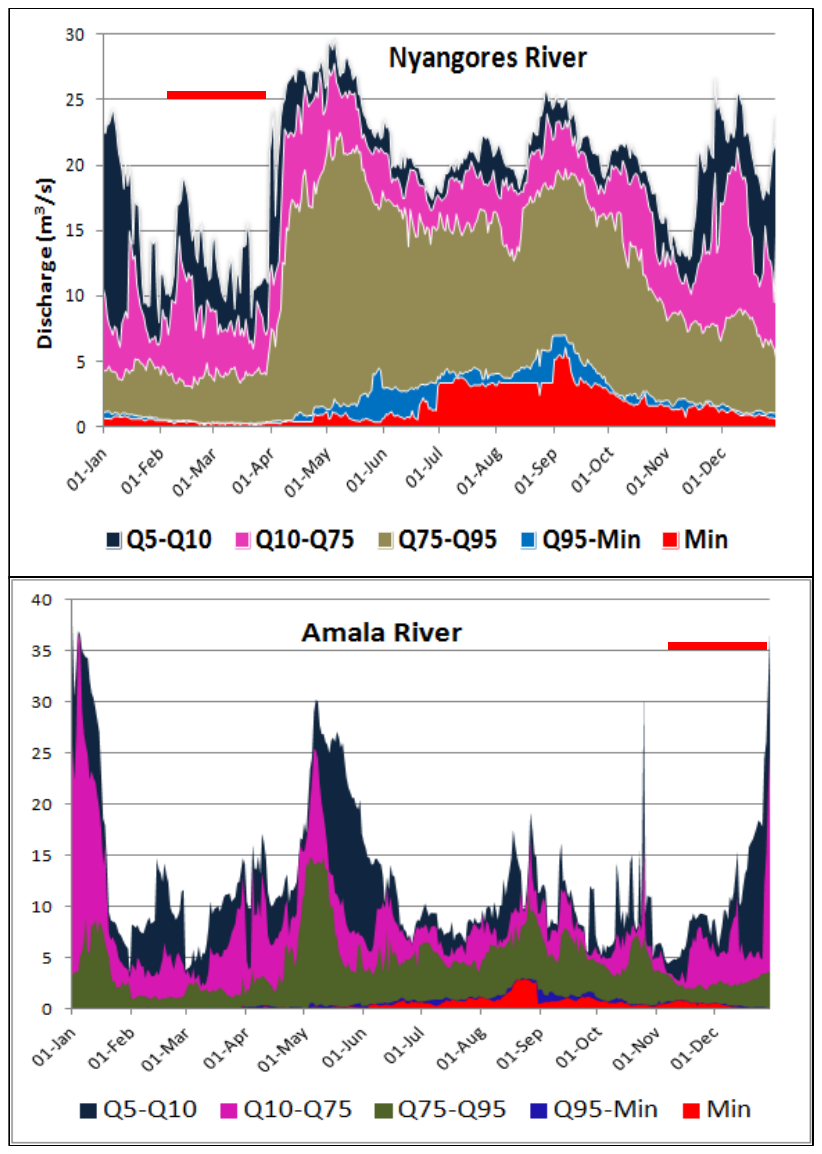
