## Supplementary Table 1 for "Livestock as vectors of organic matter and nutrient loading in aquatic ecosystems in African savannas"

**S1 Table: Bootstrap medians and 25% and 95% confidence intervals for cattle characteristics and proportions of C, N and P in dung and urine in the Mara River basin, Kenya.**

|  | **Median** | **25% CI** | **95% CI** |
| --- | --- | --- | --- |
| Time spent at watering points (minutes) | 11.5 | 10.6 | 12.5 |
| Percent defecation | 12.4 | 11.3 | 13.6 |
| Percent urination | 13.6 | 12.2 | 14.9 |
| Dung wet weight (g) | 868.5 | 749.8 | 991.1 |
| Urine volume (ml) | 0.66 | 0.53 | 0.82 |
| Percent C in urine | 14.3 | 13.5 | 15.0 |
| Percent N in urine | 10.3 | 9.7 | 11.0 |
| Percent P in urine | 0.43 | 0.33 | 0.54 |
| Percent C in dung | 32.55 | 30.20 | 34.94 |
| Percent N in dung | 1.41 | 1.23 | 1.67 |
| Percent P in dung | 0.24 | 0.28 | 0.33 |
