## Supplementary Table 2 for "Livestock as vectors of organic matter and nutrient loading in aquatic ecosystems in African savannas"

**Table S2.** Differences in density of cattle and hippopotamus, and physico-chemical characteristics (mean ± SD) across different sites in the Mara River, Kenya, grouped into five categories: Forested, Agricultural, low density (LD) livestock (mainly cattle), high density (HD) livestock, and hippopotamus (hippos). The statistics are for one-way ANOVA used to analyse significant differences in physical and chemical variables and nutrient concentrations across the five site categories in the Mara River basin, Kenya.

|  | **Site categories** | | | | | **Statistics** | |
| --- | --- | --- | --- | --- | --- | --- | --- |
| **Characteristics** | **Forested** | **Agricultural** | **Low Density Livestock** | **High Density Livestock** | **Hippos** | **F - value** | ***p* - value** |
| LMH density (individuals / km^2^) ^#^ | 9.3±4.1^a^ | 28.0±16.8^b^ | 44.8±12.5^b^ | 102.9±17.7^c^ | 111.9±10.8^c^ | 11.21 | 0.001 |
| River width (m) | 7.9±5.0^a^ | 7.1±6.1^a^ | 8.7±5.8^a^ | 6.6±3.5^a^ | 15.84±9.9^b^ | 4.35 | 0.003 |
| Water depth (m) | 0.20±0.1^a^ | 0.2±0.1^a^ | 0.2±0.2^a^ | 0.2±0.1^a^ | 1.0±0.4^b^ | 5.45 | 0.001 |
| River discharge (m^3^/s) | 0.23±0.2^a^ | 0.4±0.2^a^ | 0.5±0.4^a^ | 0.3±0.2^a^ | 17.4±6.9^b^ | 7.57 | 0.001 |
| pH (units) | 7.6±0.3^a^ | 7.6±0.3^a^ | 7.7±0.2^a^ | 7.6±0.1^a^ | 7.8±0.0^a^ | 0.40 | 0.804 |
| Temperature ⁰C | 15.8±1.8^a^ | 18.5±3.2^b^ | 20.3±2.5^b^ | 23.7±2.1^c^ | 23.9±2.2^c^ | 27.9 | 0.001 |
| Dissolved oxygen (mg/L) | 7.8±0.4^a^ | 7.0±1.1^bc^ | 6.3±0.9^c^ | 7.3±1.0^ab^ | 8.0±1.1^a^ | 7.36 | 0.001 |
| Conductivity (μS/cm) | 74.1±29.4^a^ | 103.2±50.0^a^ | 260.3±128.9^b^ | 309.0±172.8^b^ | 325.0±178.7^b^ | 18.12 | 0.001 |
| DOC (mg/L) | 2.8±1.9^a^ | 2.8±1.3^a^ | 6.4±2.9^b^ | 6.8±1.6^b^ | 5.8±2.5^b^ | 17.00 | 0.001 |
| NO_3_^-^-N (mg/L) | 0.58±0.27^ab^ | 0.8±0.4^a^ | 1.0±0.4^a^ | 0.8±0.4^b^ | 0.7±0.5^ab^ | 4.22 | 0.004 |
| NO_2_^-^-N (μg/L) | 190.7±62.4^a^ | 196.0±51.6^a^ | 294.0±60.9^a^ | 2.2±0.3^a^ | 2.1±0.5^a^ | 1.54 | 0.211 |
| NH_4_-N (μg/L) | 7.7±6.9^a^ | 97.2±64.2^a^ | 67.4±39.7^a^ | 153.9±143.3^a^ | 698.0±617.0^b^ | 14.92 | 0.001 |
| TDN (mg/L) | 0.78±0.9^a^ | 1.1±0.4^a^ | 1.3±0.6^ab^ | 0.9±0.5^ab^ | 1.4±0.6^ab^ | 3.54 | 0.041 |
| SRP (μg/L) | 9.5±8.8^a^ | 15.4±26.5^b^ | 20.2±9.5^a^ | 24.5±15.2^ab^ | 49.6±42.7^b^ | 5.51 | 0.001 |

^#^LMH – large mammalian herbivores, mainly cattle and hippopotamus
